# Tumor-microvessel on-a-chip reveals sequential intravasation cascade of cancer cell clusters

**DOI:** 10.1101/2024.02.28.582606

**Authors:** Yukinori Ikeda, Jun-ichi Suehiro, Hiroko Oshima, Sau Yee Kok, Kazuki Takahashi, Hiroyuki Sakurai, Tetsuro Watabe, Masanobu Oshima, Yukiko T. Matsunaga

**Affiliations:** Institute of Industrial Science, The University of Tokyo, 4-6-1 Komaba, Meguro-ku, Tokyo 153-8505, Japan; Department of Bioengineering, School of Engineering, The University of Tokyo, 4-6-1 Komaba, Meguro-ku, Tokyo 153-8505, Japan; Department of Pharmacology and Toxicology, Kyorin University School of Medicine, 6-20-2, Shinkawa, Mitaka, Tokyo, 181-8611, Japan; Division of Genetics, Cancer Research Institute, Kanazawa University, Kakuma-machi, Kanazawa, 920-1102, Japan; WPI Nano Life Science Institute, Kanazawa University, Kanazawa, 920-1102, Japan; Department of Biochemistry, Graduate School of Medical and Dental Sciences, Tokyo Medical and Dental University (TMDU), 1-5-45 Yushima, Bunkyo-ku, Tokyo 113-8549, Japan

## Abstract

Circulating tumor cell (CTC) clusters are often detected in blood samples of patients with high-grade tumor and are associated with tumor metastasis and poor prognosis. However, the underlying mechanisms by which CTC clusters are released from primary tumors beyond blood vessel barriers remain unclear. In this study, a three-dimensional (3D) in vitro culture system is developed to visualize tumor intravasation by positioning tumor organoids with distinct genetic backgrounds to surround microvessels. We visualized tumor intravasation in a cluster unit, including collective migration in the collagen gel, vessel co-option, and the release of CTC clusters as one of cluster invasion manners yet reported previously. In addition, our results show that both transforming growth factor-β (TGF-β) expression in tumor cells and subsequent induction of activin expression in endothelium are essential for tumor cell intravasation accompanied with endothelial-to-mesenchymal transition (EndoMT) in microvessels. Our 3D in vitro system can be used to develop therapeutic strategies for tumor metastasis by targeting CTC clusters.

## INTRODUCTION

Metastasis is the main cause of cancer-related death (*1*, *2*). Therefore, it is important to understand the blood-borne metastatic process at the molecular and cellular levels to develop therapeutic strategies and improve cancer prognosis (*3*, *4*). Recently, it has been shown that polyclonal metastasis due to tumor cell clusters is associated with higher metastatic efficiency and lower prognosis accuracy compared to the conventional metastatic process with single cell origin (*5-9*). This concept was supported by preclinical or clinical evidence, such as collective migration into stroma, vessel co-option, and the detection of circulating tumor cell (CTC) clusters from patient blood. However, it remains unclear how CTC clusters are generated from primary tumors beyond the robust endothelial barrier.

The accumulation of gene mutations in colorectal cancers is essential for acquiring metastatic potential (*10*, *11*). In particular, the combination of gene mutations, *Apc^Δ716^* (A), *Kras^G12D^*(K), *Tgfbr2^−/−^* (T), or *Trp53^R270H^* (P), contributes to collective metastasis in animal experiments using intestinal tumor model mice. For example, mouse intestinal tumor cells that have A and P gene mutations (abbreviated as AP) are invasive but nonmetastatic, whereas those with A, K, T, and P gene mutations (abbreviated as AKTP) are invasive and highly metastatic (*12*, *13*). To elucidate how gene mutations influence metastatic tumor development, in vitro culture models are required for sequential visualization of the metastatic process.

Here, we developed a three-dimensional (3D) in vitro culture system to visualize tumor intravasation by positioning tumor organoids surrounding microvessel models based on microphysiological systems technology (*14-17*). Compared with conventional in vitro cultures, which are separated by porous membranes (*18*, *19*), the hydrogel-based 3D microenvironments allow the interaction of tumor cells with the endothelial layer in a cluster unit. Moreover, unlike randomly arranged tumor organoids (*20-22*), we established distant-interaction (DI) and near-interaction (NI) models in the 3D in vitro culture system, in which the distance between tumor organoids and a microvessel is strictly controlled, to elucidate the contribution of secreted factors. Using the in vitro microvessel models and tumor cells with different genetic backgrounds (*12*, *13*), we visualized the sequential process of CTC cluster release from tumor organoids and found that transforming growth factor-β (TGF-β) is essential for tumor cell intravasation accompanying endothelial-to-mesenchymal transition (EndoMT). Our study can pave the way for the discovery of molecular targets associated with collective migration and polyclonal metastasis using drug screening and the improvement of cancer prognosis.

## RESULTS

### In vivo metastasis study and 3D in vitro tumor–microvessel models using genetically defined tumor organoids

We have previously shown that mouse intestinal tumor-derived organoids with different genotypes were established from compound mutant mice carrying combined two or more mutations, mainly *Apc^Δ716^* (A), *Kras^G12D^* (K), *Tgfbr2*^−/−^ (T), or *Trp53^R270H^* (P), in any patterns (*12, 13*). In this study, AP or AKTP organoids were transplanted to the spleen of NOD.Cg-*Prkdc^scid^Il2rg^tm1Wjl^*/SzJ (NSG) mice to observe their metastatic behavior on the extravasation from sinusoidal vessels into liver parenchyma (Fig. 1A). At day 7 after transplantation, hematoxylin and eosin-stained liver tissue sections showed that both AP and AKTP cells were detected at the lumen of the liver vessels (Fig. 1B, top). At day 14, AKTP cells invaded the liver parenchyma from the vessels, whereas AP cells remained inside the vessel lumens (Fig. 1B, bottom), indicating that compound mutation held by tumor cells might determine whether tumor cells have invaded the tissue of other organs beyond vessels during blood-borne metastasis.

**Fig. 1.**
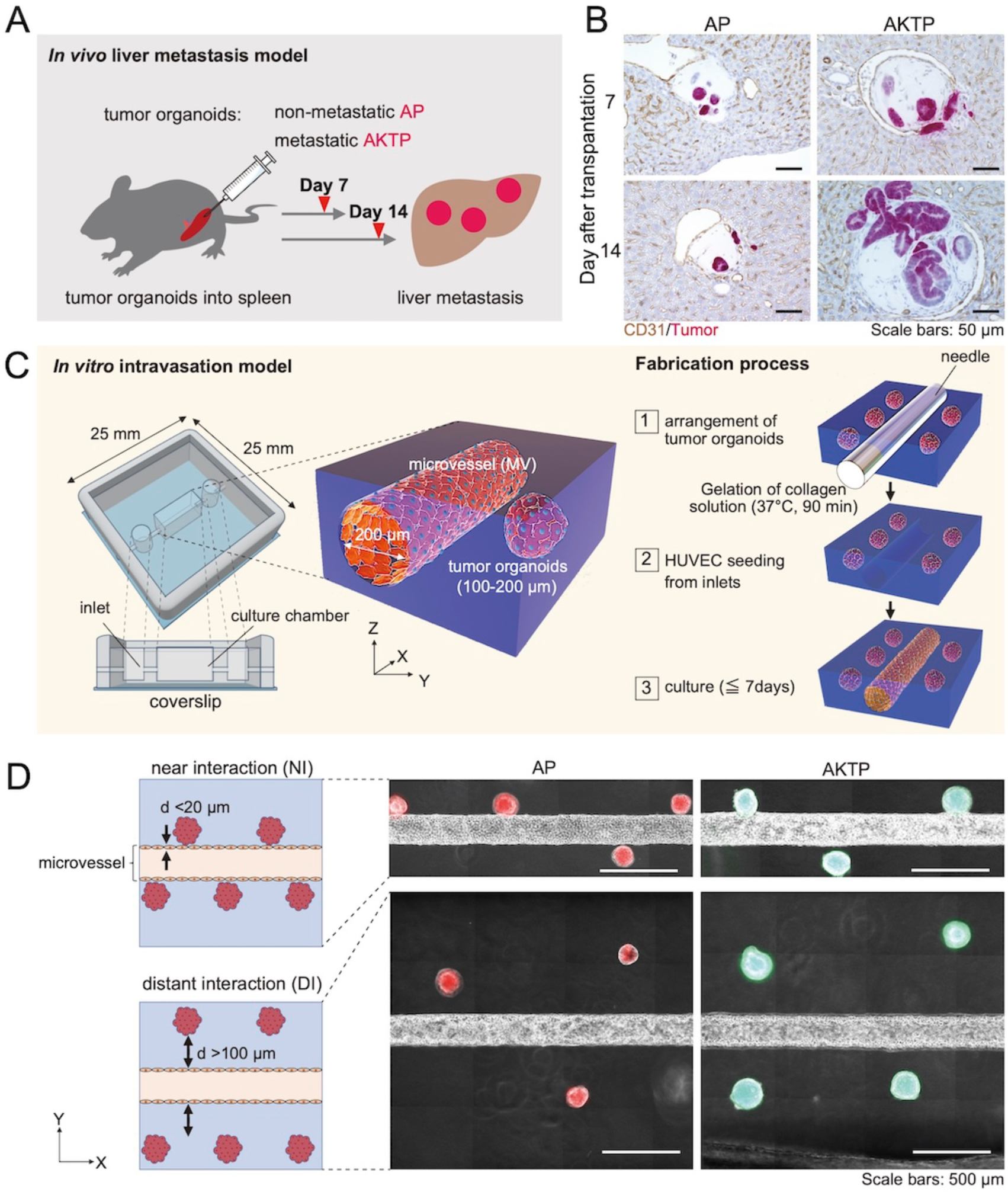
In vivo metastasis study and 3D in vitro tumor–microvessel model. (**A**) Schematic illustration of the in vivo metastasis model. Non-metastatic AP or metastatic AKTP organoids were transplanted with Matrigel into the mouse spleen. (**B**) Immunocytochemistry of liver tissues performed using antibodies against CD31 or the fluorescent protein specifically expressed on the transplanted tumor organoids. Representative tissue-sectional image of the CD31-positive sinusoidal vessels in liver with AP (left) and AKTP organoids (right). *n* = 3 biologically independent animals. (**C**) Experimental setup for the 3D in vitro tumor–microvessel model. (**D**) Schematic illustration and representative microscopic images of tumor–microvessel models with different distances (abbreviated as “d”) between the tumor organoid and the microvessel: near-interaction models (NI) and distant-interaction models (DI). tdTomato-labeled AP (red); Venus-labeled AKTP (green).

To elucidate differences in metastatic ability between AP and AKTP at the cellular level with respect to the vessel–tumor interplay, we developed 3D in vitro culture models recapitulating their spatial arrangement in the tumor microenvironment (Fig. 1C). The in vitro culture system was composed of a microvessel (200 μm in diameter) with encapsulated tumor organoids (100–200 μm in diameter) in a collagen gel. It enables spatiotemporal immunobiological analysis at the cellular level unattainable using conventional in vitro 2D cell cultures or rodent animal experiments. By manual positioning of tumor organoids, we obtained NI (<20 μm) or DI (>100 μm) models according to the distance between tumor organoids and the microvessel (Fig. 1D). NI models can be applied to observe tumor intravasation mediated by direct cell–cell interactions, whereas DI models can be used to examine metastatic potential due to secreted factors from both tumors and microvessels not accompanying direct cell–cell interactions.

### Metastatic process through direct tumor–microvessel interactions

To examine whether tumor metastatic abilities were affected by the physical contact between the tumor and endothelial cell layers, we analyzed cellular and tissue behavior using metastatic-AKTP or nonmetastatic AP organoids in the NI model. In the coculture with AKTP, time-lapse imaging revealed that AKTP organoids invaded the lumen of the microvessel at day 3 and that the release of CTC clusters into the microvessel occurred at day 5 (Fig. 2, A and B). In addition, we detected the sequential process of tumor intravasation: (i) hijacking of endothelial cell layers by AKTP cells; (ii) release of CTC clusters from AKTP organoids into the microvessel lumen (Fig. 2C, movie S1).

**Fig. 2.**
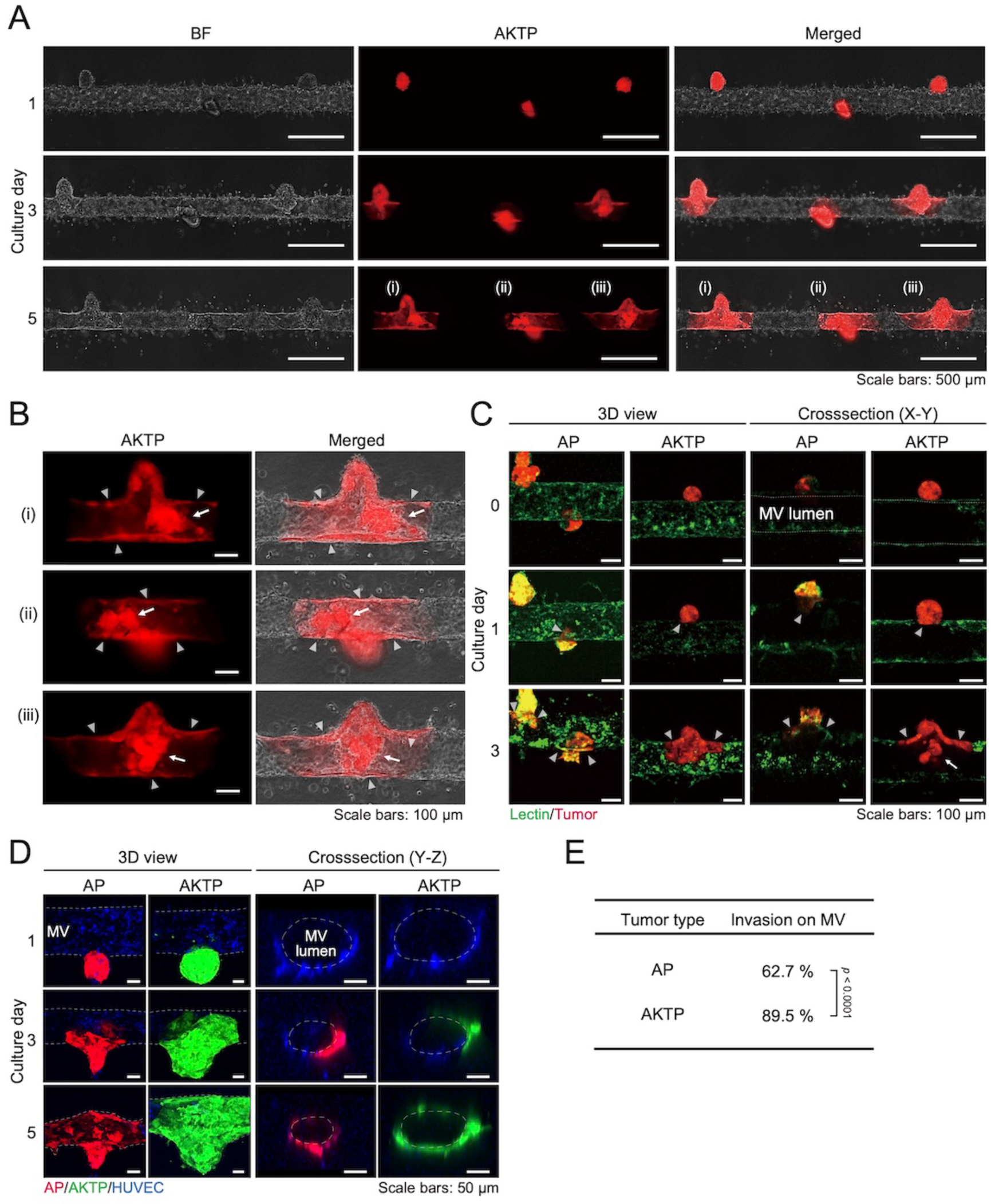
Tumor intravasation in the 3D near–interaction (NI) model: the microvessel (MV) hijacked by tumor cells and the release of CTC clusters. (**A**) Time-dependent microscopic images of intravasation by AKTP organoids (red). (**B**) Enlarged images of the intravasation region at day 5 in A. Hijacked microvessels that tumor cells invade on the microvessel inner wall (arrowheads) and the release of CTC clusters into the lumen (arrow) were observed. (**C**) Time-dependent 3D or cross-sectional CLSM images of intravasation by AP or AKTP organoids. AP or AKTP cells were labeled with tdTomato (red), and the microvessels are stained with UEA-1 (green). Tumor invasion on the microvessel inner wall and the release of CTC clusters into the lumen were indicated by arrowheads and arrows, respectively. The microvessel layer (green) at day 0 was indicated by dotted line (movie S1 and S2). (**D**) Time-dependent CLSM images of the microvessel hijacked by tdTomato-labeled AP (red) or Venus-labeled AKTP (green) organoids. Microvessels were stained with CellTracker violet BMQC (blue). The outline of the microvessel and the area hijacked by AP or AKTP organoids was indicated by the dotted line. (**E**) Ratio of the AP- or AKTP- invasion area on the microvessel at day 5. (*n* = 4-5 for biological chip replicates). The data were analyzed by two-sided chi-square and Fisher’s exact test to calculate the significance of differences. *p* values are provided.

Although both AKTP and AP invaded the microvessel layer, the release of CTC clusters by non-metastatic AP cells was hardly detected (Fig. 2C, movie S2). The confocal laser scanning microscopy (CLSM) analysis revealed that the tumor invasion area on the microvessel lumen of both AKTP and AP gradually increased with time for 3 days (Fig. 2, C, arrowheads, and D). This process is known as *vessel co-option*, which is a non-angiogenic process through which tumor cells utilize pre-existing tissue blood vessels to support tumor growth, survival, and metastasis (*20*, *22*, *23*). Interestingly, the ratio of the invasive area on the microvessel lumen was 62.7% for AP and 89.5% for AKTP (Fig. 2E), supporting previous reports showing that AKTP is more invasive than AP (*12*, *13*). Taken together, the NI model can mimic sequential metastasis, such as tumor cell intravasation on the microvessel lumen and subsequent CTC cluster release with direct cell–cell interactions between tumor organoids and vessels.

### Metastatic process through tumor–microvessel secretion factors

To investigate the effects of secretion factors on tumor invasiveness, we analyzed the dynamics of cell behavior using the DI model. In the coculture between the tumor and the microvessel, AP organoids did not show migration toward the microvessel and remained in the primary position even after the 3-day culture period. In contrast, AKTP organoids exhibited invadopodium formation, detachment from the primary tumor, and collective cell migration toward the microvessel, which can be associated with tumor invasiveness and metastasis (Fig. 3A, movie S3 and S4). In addition, AKTP can induce endothelial cell migration toward the organoids (Fig. 3A, circle). The ratio of invadopodium formation on tumor organoids in the coculture with the microvessel at day 3 increased more than four times compared to that in the tumor monoculture (Fig. 3, B and C).

**Fig. 3.**
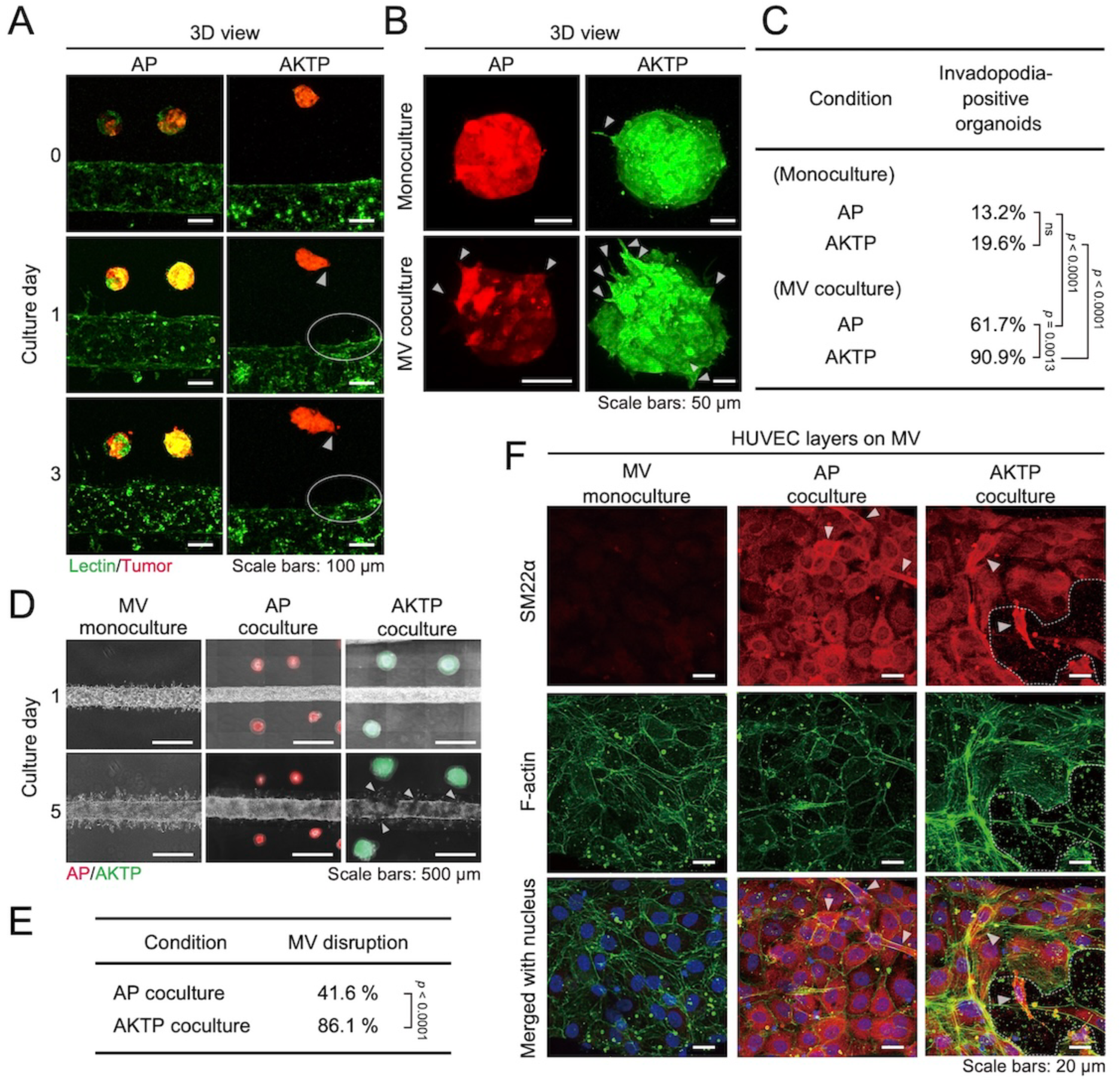
3D distant–interaction (DI) models show metastasis-related phenotypic changes through cell–cell crosstalk by secreted factors. (**A**) Representative 3D CLSM images of time-dependent interaction between AP or AKTP and microvessels. AP or AKTP cells are labeled with tdTomato (red) and microvessels are stained with UEA-1 (green). The migration of AKTP organoids or angiogenic sprouts were indicated by the arrowheads or circles, respectively. (**B**) Representative 3D CLSM images of AP or AKTP organoids in monoculture or in coculture with microvessel at day 3. The formation of invadopodia is indicated by the arrowheads. AP or AKTP cells were labeled with tdTomato (red) or Venus (green). (**C**) Ratio of AP or AKTP organoids with invadopodia (>20 µm at length) in monoculture or coculture with microvessels at day 3. (*n* = 3-4 for biological chip replicates). (**D**) Time-dependent microscopic images of microvessels with disrupted regions near tumor organoids at day 5. AP or AKTP were labeled with tdTomato (red) or Venus (green), respectively. The disrupted regions of microvessel edges were indicated by the arrowheads. (**E**) Ratio of disrupted microvessel regions nearby tumor organoids in tumor–microvessel models at day 5. (**F**) Immunofluorescent CLSM images of the HUVEC layer on the tumor–microvessel models at day 5. SM22α-positive HUVECs or disrupted area on the HUVEC layer were indicated by the arrowheads or the dotted line, respectively. (*n* = 8-9 for biological chip replicates). The data in C and E were analyzed by two-sided chi-square and Fisher’s exact test to calculate the significance of differences. *p* values are provided. ns not significant.

Interestingly, endothelial cell layers on the microvessel were partially disrupted near tumor organoids at day 5 in the DI model. These endothelial cell disruptions were more often observed in the coculture with AKTP (86.1%) rather than that with AP (41.6%) (Fig. 3, D and E). During tumor metastasis to distant organs, it has been reported that vascular integrity was reduced to facilitate tumor cell intravasation and extravasation. This endothelial barrier disruption is induced by endothelial-to-mesenchymal transition (EndoMT; also called EndMT), which plays a role in tumor progression and metastasis (*24*, *25*). During EndoMT, endothelial cells lose their characteristics, such as strong cell–cell contact and the expression of endothelial cell-specific markers, including vascular endothelial growth factor receptor 2 (VEGFR2, KDR) and tunica interna endothelial cell kinase 2 (Tie2), and acquire mesenchymal phenotypes, such as stress fiber formation and the expression of mesenchymal cell markers, including smooth muscle 22α (SM22α), α-smooth muscle actin (αSMA), and fibronectin. Therefore, we examined mesenchymal marker expression and stress fiber formation in endothelium. SM22α, a marker of mesenchymal cells, and phalloidin staining were significantly upregulated by the coculture with AKTP but not the endothelial monoculture (Fig. 3F).

Taken together, the coexistence of tumor organoids and the microvessel can alter endothelial characteristics toward a metastatic phenotype, such as EndoMT, leading to a malignancy of the tumor microenvironment, which might be a prerequisite for tumor metastasis.

### Morphological and molecular changes underlying tumor–microvessel interactions

To understand the molecular mechanisms underlying intercellular signal transduction between tumor organoids and the microvessel, we used a semi-3D coculture model (Fig. 4, A and B). The semi-3D coculture model facilitates the observation of cell morphologies in both tumor organoids and endothelial layers and allows qPCR gene expression analysis as it provides a simpler format than the 3D system.

**Fig. 4.**
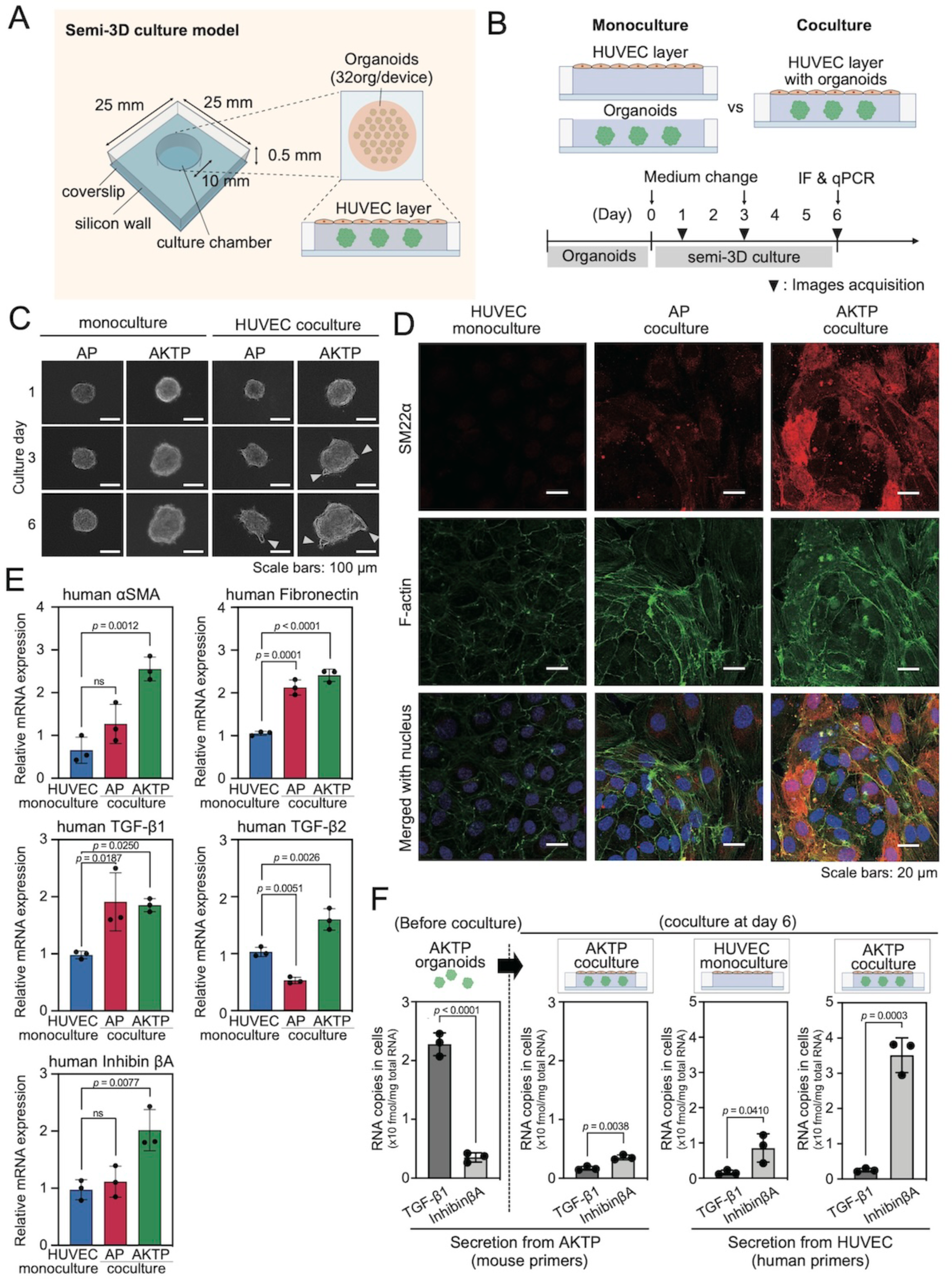
TGF-β family ligands are key mediators of malignant crosstalk between tumor and endothelial cells. (**A**) Experimental setup for the in vitro semi-3D culture model of gene expression in tumor organoids and endothelial cells. (**B**) Schematic illustration of the experimental condition and time course in the semi-3D study. (**C**) Representative time-dependent phase-contrast images of AP or AKTP organoids under monoculture or coculture with the endothelial layer. Invadopodia were indicated by arrowheads. (**D**) Representative CLSM images showing the immunostained HUVEC layer in monoculture or cocultured with AP or AKTP organoids at day 6 after coculture. (**E**) Relative mRNA levels of mesenchymal markers and TGF-β family ligands in HUVECs cultured with or without AP or AKTP organoids at day 6 after coculture. Inhibin βA indicates subunits of activin. (*n* = 3 for biological chip replicates) (**F**) Determination of mRNA copies of TGF-β1 and Inhibin βA (subunits of activin) in HUVEC and AKTP cells at day 6 before or after coculture. (*n* = 3 for biological chip replicates) The data in E and F are presented as the mean ± s.d. Significant differences were analyzed by one-way ANOVA in E or two-tailed unpaired Student’s *t*-test in F, respectively. *p* values are provided, ns not significant.

Similar to the results observed by the 3D model, coculture in the semi-3D model showed metastatic and EndoMT behaviors at morphological and protein expression levels. Tumors showed metastatic invadopodium formation when cocultured with endothelial layers; endothelial cells showed the expression of SM22α as an EndoMT marker and the disruption of endothelial layers (Fig. 4, C and D). These phenomena were further enhanced in AKTP than in AP in the coculture system. Further, quantitative PCR analysis revealed mesenchymal markers on the endothelial layers: fibronectin, and αSMA, were significantly upregulated when cocultured with tumor organoids (Fig. 4E).

TGF-β family members, including TGF-β1, TGF-β2, TGF-β3, and activin A: a subunit of Inhibin βA (hereafter termed activin), are involved in various stages of tumor progression and metastasis, including epithelial-to-mesenchymal transition (EMT) in epithelial tumor cells and EndoMT (*26*). TGF-βs and activin activate and transduce their signals via specific receptor complexes, including type I (TβRI and ActRI) and type II (TβRII and ActRII) receptors for TGF-βs and activin, respectively. Quantitative PCR analysis also revealed TGF-β family on the endothelial layers: TGF-β1, TGF-β2, and activin were significantly upregulated when cocultured with AKTP organoids (Fig. 4E).

Given that TGF-βs and activin may contribute to invadopodium formation and EndoMT, we investigated differences in gene expression among ligands for the TGF-β family by calculating RNA copies in cells where RNA expression derived from tumor cells or human umbilical vein endothelial cells (HUVECs) can be distinguished by primers specific for mice or human genes, respectively. Quantitative PCR showed that TGF-β1 was originally expressed more than Inhibin βA in AKTP cells before the coculture with endothelium (Fig. 4F). On the other hand, activin expression was upregulated four folds in the endothelial cell layer at day 6 after the coculture with AKTP compared with the monoculture (Fig. 4F). Notably, only activin but not TGF-β1 may contribute to invadopodium formation in the coculture shown in Fig. 4C because AKTP cells have no functional receptors for TGFβ1-β3 due to *Tgfbr2* deletion.

Taken together, the results obtained from the semi-3D coculture model suggested that the coculture with AKTP cells promoted invadopodium formation and EndoMT, which might be mediated by the original expression of TGF-β1 in AKTP cells and the subsequent secretion of activin from mesenchymal-like endothelial cells.

### Role of TGF-β1, -β2, and -β3 in tumor–microvessel interactions

TGF-β1, TGF-β2, and activin have been shown to play a pivotal role in intestinal tumor progression and have malignant effects on the tumor microenvironment (*13*, *26*, *27*). From the results shown in Figs. 1 to 4, we found that AKTP was more invasive and promoted EndoMT more vigorously than AP. Therefore, we focused on the AKTP coculture with and without the endothelial cell layer to examine the effects of TGF-β family ligands on tumor aggressiveness. To investigate the role of TGF-β1 and TGF-β2 in facilitating tumor invasion and EndoMT, an Fc-chimeric protein containing the extracellular domains of both TGF-β type I and II receptors (TβRI-TβRII-Fc) was introduced into the semi-3D model (Fig. 5, A and B), which has been demonstrated to inhibit the TGF-β signal, effectively and specifically neutralizing TGF-β1, TGF-β2, and TGF-β3 (*28*).

**Fig. 5.**
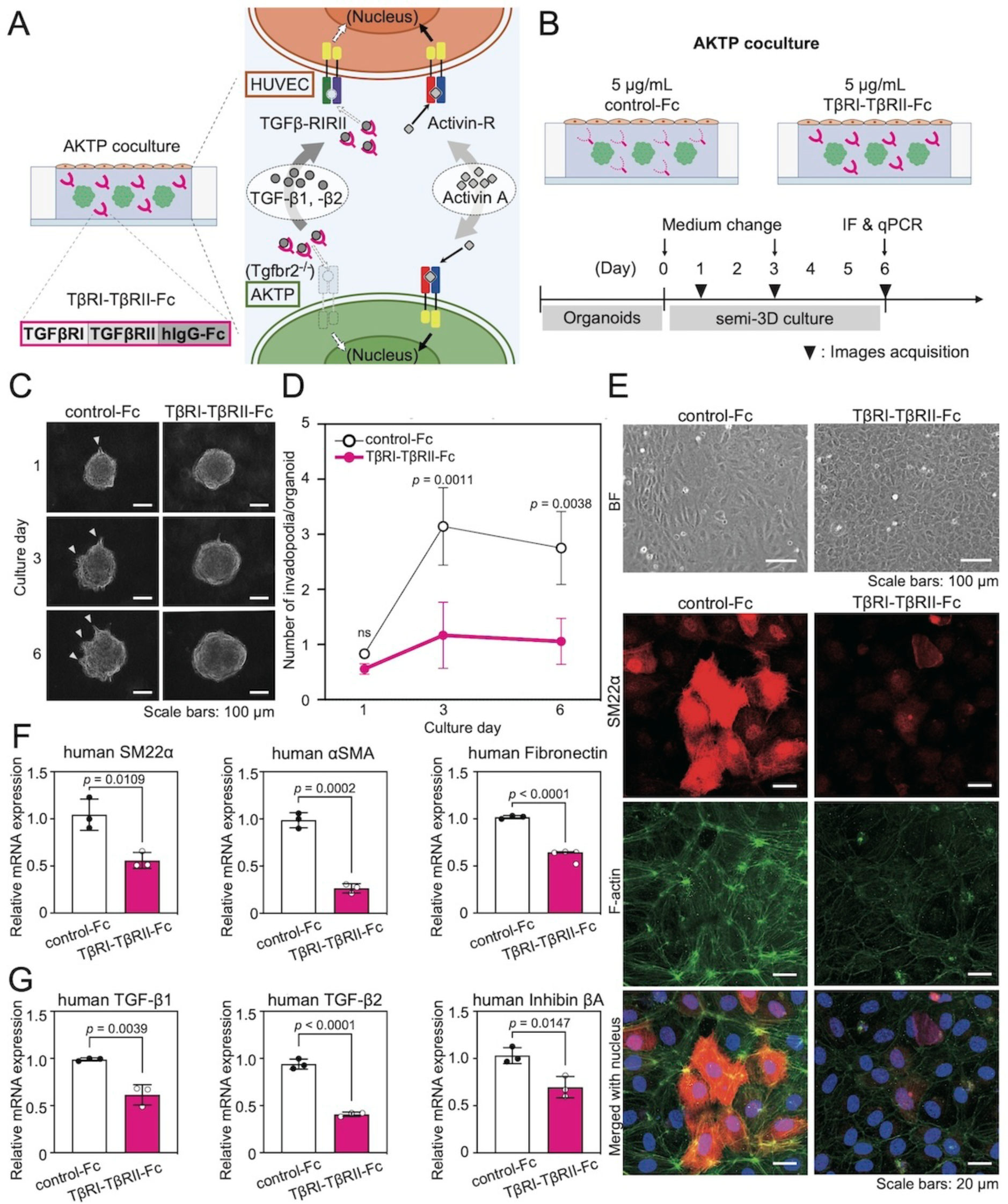
TGF-β1 and its isoforms are potent initiators of cell–cell crosstalk between tumor and endothelial cells. (**A**) Schematic illustration of the concept of in vitro semi-3D experiment treated with the Fc-chimeric protein. TβRI-TβRII-Fc specifically capture TGF-β1, -β2, and -β3 secreted by the HUVEC layer or AKTP organoids. (**B**) Schematic illustration of in vitro semi-3D experiment. The Fc region of an IgG (control-Fc) was used as a control. (**C**) Representative time-dependent phase-contrast images of AKTP organoids with the HUVEC layer treated with TβRI-TβRII-Fc or control-Fc. Invadopodia are indicated by the arrowheads. (**D**) Quantification of invadopodium formation on AKTP organoids under coculture with HUVEC in the treatment of TβRI-TβRII-Fc or control-Fc. (*n* = 3 for biological chip replicates) (**E**) Representative phase-contrast images (top) and CLSM images (bottom) showing the immunostained HUVEC layer with AKTP organoids at day 6 after treated with TβRI-TβRII-Fc or control-Fc. (**F**) Relative mRNA levels of mesenchymal markers in HUVEC with AKTP organoids at day 6 after treated with TβRI-TβRII-Fc or control-Fc. (*n* = 3 for biological chip replicates) (**G**) Relative mRNA levels of TGF-β family ligands in HUVEC at day 6 after treated with TβRI-TβRII-Fc or control-Fc. Inhibin βA indicates subunits of activin. (*n* = 3 for biological chip replicates) The data in D, F and G are presented as the mean ± s.d.. Significant differences between control and treated condition were analyzed by two-way ANOVA in D or two-tailed unpaired Student’s *t*-test in F and G, respectively. *p* values are provided.

The microscopic image analysis showed that invadopodium formation on AKTP was reduced to 30% under the TβRI-TβRII-Fc treatment compared with control-Fc (Fig. 5, C and D). TβRI-TβRII-Fc also preserved the normal pavement shape in endothelium (Fig. 5E). The quantitative PCR analysis showed that SM22α, αSMA, and fibronectin expression remarkably decreased in endothelium upon the TβRI-TβRII-Fc treatment compared with control-Fc (Fig. 5F). Moreover, TGF-β family genes were prominently downregulated in the endothelial layer (Fig. 5G). These results suggested that the specific inhibition of TGF-β1-β3 signaling suppressed the gene expression of the mesenchymal marker and TGF-β ligands in endothelium and further attenuated both EndoMT and AKTP invasion.

### Understanding the role of activin in tumor–microvessel interactions

To elucidate the role of activin in coculture with AKTP and endothelial layer, follistatin, a well-known activin inhibitor (*29*), was introduced into the semi-3D model (Fig. 6, A and B). The morphological analysis conducted by the microscopic image showed that invadopodium formation on AKTP was significantly reduced to less than 30% under the treatment of follistatin compared with the control, which was similar to the effects of TβRI-TβRII-Fc (Fig. 6, C and D). On the other hand, follistatin failed to suppress the disruption of the endothelial layer and reversed the upregulation of mesenchymal markers in endothelial cells (Fig. 6E). The gene expression analysis using qPCR showed that αSMA was significantly suppressed by follistatin, whereas SM22α and fibronectin remained unchanged compared with the control (Fig. 6F). With respect to TGF-β family ligands, TGF-β2 and Inhibin βA expressions were slightly downregulated in endothelium (Fig. 6G). Taken together, follistatin suppressed AKTP invasiveness but had a partial effect on EndoMT and the expression of TGF-β family ligands in the coculture.

**Fig. 6.**
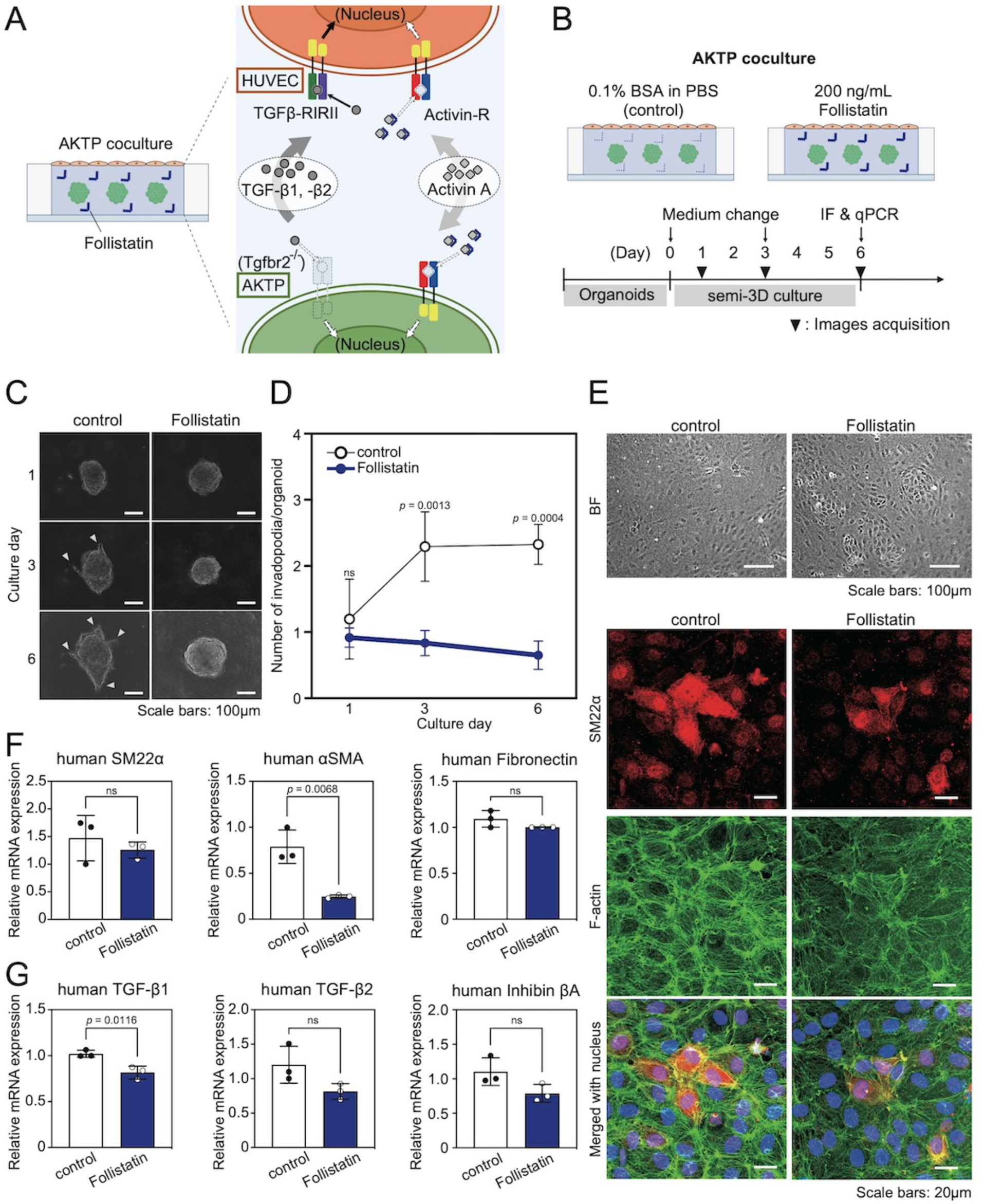
Activin secretion from TGF-β-activated endothelial cells plays a pivotal role for AKTP invasiveness. (**A**) Schematic illustration of the concept of in vitro semi-3D experiment treated with follistatin. Follistatin neutralizes activin secreted by HUVEC or AKTP organoids. (**B**) Schematic illustration of the in vitro semi-3D experiment. 0.1% BSA in PBS as a vehicle was added to the control. Image acquisition and expression analysis were performed at the indicated time points. (**C**) Representative time-dependent phase-contrast images of AKTP organoids with the HUVEC layer treated with follistatin or its vehicle alone. Invadopodia are indicated by the arrowheads. (**D**) Quantification of invadopodium formation on AKTP organoids under coculture with HUVEC treated without (control) or with follistatin. (*n* = 3 for biological chip replicates) (**E**) Representative phase-contrast images (top) and CLSM images (bottom) of the immunostained HUVEC layer with AKTP organoids at day 6 after treated without (control) or with follistatin. (**F**) Relative mRNA levels of mesenchymal markers in HUVECs with AKTP organoids at day 6 after treated without (control) or with follistatin. (*n* = 3 for biological chip replicates) (**G**) Relative mRNA levels of TGF-β family ligands in HUVEC at day 6 after treated without (control) or with follistatin. Inhibin βA indicates subunits of activin. (*n* = 3 for biological chip replicates) The data in D, F and G are presented as the mean ± s.d.. Significant differences between control and treated condition were analyzed by two-way ANOVA in D or two-tailed unpaired Student’s *t*-test in F and G, respectively. *p* values are provided. ns not significant.

## DISCUSSION

We developed a 3D in vitro intravasation model with the arranged distance between tumor organoids and the microvessel to understand molecular mechanisms that mediated distinct malignancy among tumor cells with different genotypes at the cellular and molecular levels (Fig. 1). In addition, we visualized the time-dependent behavior of AKTP tumor organoids during intravasation, including the disruption of the endothelial cell layer, hijacking of the microvessel lumen, and release of tumor cell clusters as a model of CTC clusters (Fig. 2 to 4). Moreover, we found that the malignancy crosstalk between endothelial and tumor cells was mediated by TGF-β1, -β2, and activin secretion from each cell, which promoted EndoMT and tumor invasion (Fig. 4 to 6).

Our 3D in vitro system offers three advantages to evaluate tumor metastasis compared to conventional in vitro systems (*20*, *21*). First, by manipulating the distance between tumor cells and microvessels in collagen gel, we achieved contact (NI model) and non-contact (DI model) between tumor organoids and microvessels. Second, we introduced colorectal tumor organoids with distinct genetic backgrounds and different in vivo metastatic potential into this device. Finally, this unique concept and device fabrication enabled us to reproduce the early phase of metastatic process, such as collective migration from the primary tumors and intravasation, and compare the morphological changes between tumors with different genetic backgrounds, AP and AKTP. Although the concept of polyclonal metastasis has been confirmed by in vivo studies, it has not yet been fully understood how CTC clusters are generated, namely, cancer cells aggregate inside vessels or cell clusters invade it. In our device model, we found the release of cell clusters from AKTP organoids through the sequential metastatic and EndoMT process including invadopodium formation, microvessel layer disruption, and invasion into the microvessel lumen. Such unique intravasation process has rarely found when AP organoids contacted to vessels. Taken together, these results suggest the novel mechanism of malignant cancer cell clusters, which subsequently circulate as CTC clusters. The present results also suggest that nonmetastatic cells cannot generate CTC clusters even if they can intravasate.

In vivo invadopodium formation and vessel co-option have been widely reported for breast or pancreatic tumors as they are common among the majority of tumor types (*20*, *22*, *23*, *30-32*). However, their role in the promotion of metastasis and molecular mechanisms underlying the metastatic process remained unclear. Tumor cells cannot invade only by themselves but through interactions with surrounding nontumor cells (*27*, *33*, *34*). We previously reported that α-SMA-positive myofibroblasts and abnormal deposition of collagen fibers were observed at the invasion area in the intestinal submucosal region of intestinal tumor model mice with various combinations of gene mutation (e.g., AP or AT mice), suggesting that fibroblasts in stroma can facilitate cancer invasion (*12*). The tumor microenvironment comprises vascular endothelial cells and these immune or stroma cells, which tend to secrete inflammatory cytokine and growth factors. In general, normal endothelial cells exhibited a static pavement shape, whereas endothelial cells activated by secreted factors from stroma cells or tumor cells were altered to a mesenchymal-like phenotype and contributed to disease progression. Previous reports suggested that fibroblasts in breast cancer organoids have an essential role in angiogenesis in the organoids from nearby endothelial layers, which is an example of malignancy mediated by activated endothelium often seen in the tumor microenvironment (*35*). With respect to the cancer–stromal cell crosstalk in tumor progression, hepatic stellate cells activated by cancer-derived TGF-β are considered as a key mediator for cancer extravasation from sinusoidal vessels (*13*, *36*).

The TGF-β family comprises three isoforms: TGF-β1, -β2, and -β3, which are agonists for a heterodimer receptor of TβRI and TβRII. TGFβ-mediated intracellular signaling contributes to tumor progression by eliciting EMT or tumor angiogenesis via EndoMT. We have previously reported that the Fc-chimeric protein combined with TGF-β receptors inhibited tumor growth and angiogenesis in vivo (*28*). Given that the TGF-β family can facilitate AKTP invasiveness via EndoMT, we introduced TβRI-TβRII-Fc into our in vitro culture system, which inhibited tumor invasion and EndoMT, suggesting that TGF-β ligands was essential for malignancy in AKTP. We found that TGF-β induced the production of activin in lymphatic endothelial cells and play a vital role in EndoMT via cytokines (*37*). Given that activin derived from endothelium can affect AKTP, we introduced follistatin into our in vitro culture system and found that activin contributed to AKTP invasiveness but did not affect EndoMT, which was a critical modulator of the malignant process in tumors.

The Cancer Genome Atlas has shown several genetic mutations and deletions associated with colorectal tumor development and its malignant transformation (*38*); TGFBR2, a TGF-β family receptor found in this project, showed potential to promote malignant transformation and metastasis via TGF-β signaling. This contradicts the in vivo study conducted by our group concluding that AKTP (mutated Tgfbr2) has a higher metastatic potential than AP (intact Tgfbr2). To resolve this controversy, we propose a multistep malignant loop model via cell-to-cell crosstalk: i) vascular endothelial cells are activated by TGF-β family ligands from tumors and exhibit EndoMT, which is a progressive state in endothelium facilitating tumor invasion; ii) invadopodium formation on tumor cells is elicited by activin secreted by endothelial cells (Fig. 7). To the best of our knowledge, these metastatic-related behaviors have been quantitatively determined in vitro and in vivo among various tumor cell types for the first time. In addition, the role of TGF-β family ligands in tumor–endothelial malignant crosstalk for metastasis has been revealed. This in vitro intravasation model is expected to be applied to order-made medicine, such as prognosis prediction and drug screening for metastasis, by reconstituting the tumor microenvironment.

**Fig. 7.**
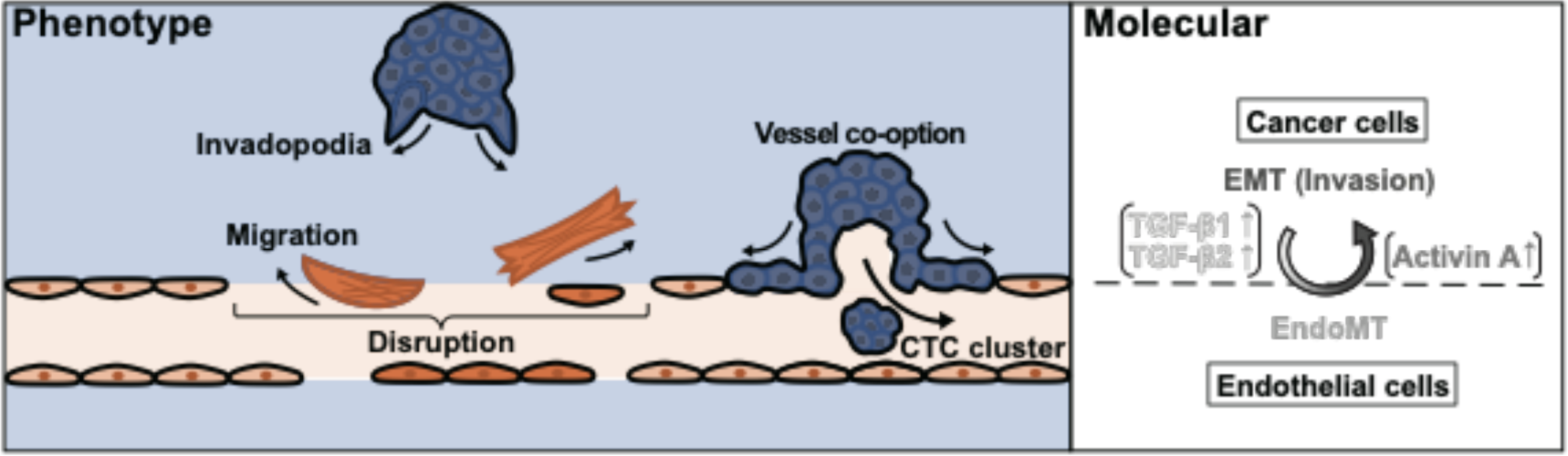
Schematic illustration of the tumor intravasation model mediated by tumor-endothelial interaction.

In summary, we visualized tumor intravasation in a cluster unit, including collective migration in the collagen gel, vessel co-option, and the release of CTC clusters as one of cluster invasion manners yet reported previously by developing novel in vitro culture systems. TGF-β family ligands were found to be key mediators of tumor invasiveness beyond the endothelium barrier by comparing the cell function among different tumor cell types with distinct genetic backgrounds. This study should be expected to contribute to developing effective strategies to halt tumor metastasis.

## MATERIALS AND METHODS

### Endothelial cells

Primary human umbilical vein endothelial cells (HUVEC; Lonza, Basel, Switzerland) were cultured in Endothelial Cell Growth Medium-2 BulletKit (EGM-2; Lonza) at 37°C in a humidified atmosphere of 5% CO_2_. The cells were used between passages 3 to 4.

### Intestinal tumor-derived cells

Mouse intestinal tumor-derived cells were used as previously described (*12*, *13*). Briefly, cell lines of AP, ATP, and AKTP were developed from intestinal tumors of *Apc^Δ716^ Trp53^R270H^*, *Apc^Δ716^ Tgfbr2^−/−^ Trp53^R270H^*, and *Apc^Δ716^ Kras^G12D^ Tgfbr2^−/−^ Trp53^R270H^*mouse intestinal tumors, respectively. AP and AKTP cells were labeled with fluorescent proteins, tdTomato (AP-tdt, AKTP-tdt) and Venus (AKTP-v), respectively. cDNAs of tdTomato and Venus were subcloned to the pPB-CAG-IP PiggyBac transposon expression vector and cotransfected with the transposase expression vector to cells using lipofectamine (Thermo Fisher Scientific, Waltham, MA, USA). Consequently, transfected clones were selected by drug selection with 1 μg/ml of puromycin (InvivoGen, San Diego, CA, USA). Mycoplasma testing was performed using an indirect immunofluorescence test. These tumor cells were cultured on dishes with the tumor 2D medium (Advanced DMEM/F-12 medium; Gibco, Thermo Fisher Scientific, Waltham, MA, USA) supplemented with 10% fetal bovine serum (FBS), 5% penicillin–streptomycin solution (FUJIFILM Wako, Osaka, Japan), 5 μM ALK inhibitor (A83-01, Sigma-Aldrich, Saint Louis, MO, USA), 5 μM GSK3 inhibitor (CHIR99021, Sigma-Aldrich), and 10 μM ROCK inhibitor (Y27632, FUJIFILM Wako). The cells were used between passages 3 to 20.

### Formation of tumor organoid

Tumor cells were suspended in the 2D media at 5.0 × 10^3^ cells/ml and then seeded into the ultralow attachment 384-well round-bottom plates (Sumitomo Bakelite, Tokyo, Japan) at 100 cells/well to form organoids. The cells were cultured for approximately 60 h, collected into a 1.5-mL tube, and centrifuged three times at 6000 rpm. The supernatant was carefully removed, and the organoids were suspended in 2.4 mg/mL neutralized type I collagen solution (Cellmatrix type I-A; Nitta Gelatin, Osaka, Japan) for further 3D or semi-3D model.

### In vivo metastasis study: transplantation of tumor cells

NSG mice were purchased from Charles River, Yokohama, Japan). The mice were housed in specific-pathogen-free conditions with a 12-h light:dark cycle at 23°C ± 2°C with a relative humidity of 50% ± 20% and given ad libitum food and water for the duration of the study. For the chronological analysis at the early stage of colonization, 5 × 10^5^ AKTP-v or 1 × 10^6^ AP-tdt cells were injected into the NSG mouse spleen with 25 μl of Matrigel (Corning, Corning, NY, USA). Liver tissues were examined histologically at day 7 and 14 after transplantation. All animal experiments were performed with the protocol approved by the Committee on Animal Experimentation of Kanazawa University (AP-204139).

The liver tissues were fixed in 4% paraformaldehyde (PFA), embedded with paraffin, and sectioned at a thickness of 4 μm. For immunohistochemistry, antibodies against GFP (#2956, Cell Signaling Technology, Danvers, MA, USA), RFP (#600-401-379, Rockland Immunochemicals INC., Limerick, PA, USA), and CD31 (#DIA-310, Clone SZ31; Dianova, Hamburg, Germany) were used as primary antibodies. The antibodies for GFP or RFP were used to detect Venus or tdTomato-labeled cells, respectively. The alkaline phosphatase-conjugated antibody against rabbit IgG (ImmPRESS AP reagent KIT, #MP-5401; Vector Laboratories, Newark, CA, USA) was used as a secondary antibody. For double-labeling immunohistochemistry, CD31 or fluorescent proteins were stained with VECTOR ImmPACT DAB peroxidase substrate kit (Vector Laboratories) or VECTOR red alkaline phosphatase substrate kit (Vector Laboratories), respectively.

### 3D tumor-microvessel model

To prepare a 3D microvessel with tumor organoids in the collagen gel, in-house polydimethylsiloxane (PDMS)-based chips (25 mm × 25 mm × 5 mm: width × length × height) were used as described previously (*16*, *17*). Tumor organoids mixed with a neutralized collagen solution (2.4 mg/mL) were added into the central chamber of the device. The BSA-coated acupuncture needle (200 μm in diameter; Seirin, Shizuoka, Japan) was inserted into the chip to form a lumen structure in the collagen gel. After manual positioning of the tumor organoid using the edge-cut BSA-coated needle (300 μm in diameter) under the microscope, the devices were turned upside down and incubated under a humidified atmosphere of 5% CO_2_ at 37°C for 60 min. Both collagen gels remained in the side chambers, and inserted needles were removed. To form microvessels, HUVECs were seeded into the collagen gel channel with a media containing 3% dextran (Leuconostoc spp., Mr 450 000–650000, Sigma-Aldrich) at a density of 1 × 10^7^ cells/mL. After microvessel formation for 10 min, media were replaced with EGM-2 for subsequent coculture with the microvessel and tumor organoids in the collagen gels and changed every 1-2 days.

### Semi-3D coculture model

The HUVEC layer on the top of the collagen gel encapsulating tumor organoids, so-called semi-3D coculture model, were prepared as follows. The in-house culture well chamber made of silicone sheets (diameter: 10 mm, height: 0.5 mm) on a glass coverslip (25 mm x 25 mm) were used only in this model. Briefly, tumor organoids mixed with an ice-cold neutralized collagen solution were added into wells at 30 organoids/well. The position of each organoid at X- and Y-axis was adjusted manually by the edge-cut BSA-coated needle under a stereomicroscope as described above. After positioning each organoid, the whole devices were turned upside down and incubated at 37°C for 45 min. HUVECs in EGM-2 containing 3% dextran at a density of 1 × 10^7^ cells/mL were added onto the surface of collagen gels and incubated at 37°C for 5 min. Cells were cultured in 2 mL of EGM-2 medium under a humidified atmosphere of 5% CO_2_ at 37°C for up to 6 days.

### Chemical inhibition study for TGF-β family members

For TGF-β family (TGF-β1, TGF-β2, and TGF-β3) inhibition, 5 µg/mL recombinant Fc-chimeric TGF-β receptor containing both TβRI and TβRII (TβRI-TβRII-Fc) or control-Fc (human IgG-Fc) was introduced to the semi-3D coculture model. TβRI-TβRII-Fc was previously shown to trap TGF-β1, -β2, and -β3 ligands specifically as a decoy receptor (*28*). For activin inhibition, we introduced 200 ng/mL follistatin (4889-FN-025, R&D Systems), an endogenous antagonist for activin (*29*), or 0.1% BSA in PBS as a vehicle. Inhibitors were added to collagen gels and the culture medium (day 0). Cells were cultured in 2 mL of EGM-2 medium under a humidified atmosphere of 5% CO_2_ at 37°C until day 6 by replacing the medium containing inhibitors at day 3. Microscopic observation under an inverted microscope was performed every 3 days, and samples were fixed for immunostaining at day 6.

### RNA isolation and quantitative RT-PCR

Total RNAs of the semi-3D culture model samples were extracted by ISOGEN (Nippon Gene, Tokyo, Japan) (*n* = 3, biologically independent). cDNA was synthesized from 0.5 or 1 μg/10 μL RNA using ReverTra Ace qPCR RT Master Mix (TOYOBO, Osaka, Japan). Real-time PCR was performed using a StepOnePlus real-time PCR system (Applied Biosystems, Thermo Fisher Scientific) or Applied Biosystems 7300 Real-Time PCR System (Applied Biosystems, Thermo Fisher Scientific) with Thunderbird SYBR qPCR Mix (TOYOBO) according to the manufacturer’s protocol. For relative quantification of human gene expression, the target gene expressions were normalized to the internal control gene, human cyclophilin A (PPIA). The level of mRNA of each gene in the semi-3D culture model was applied for the standard curve compared to the control (relative expression = 1). For determining RNA copies in cells, the titration curve was generated by successively diluting the concentration of TGF-β1 or Inhibin A PCR amplicon inserted into the TOPO cloning vector (Thermo Fisher Scientific). The sequences of primer pairs are listed in Table S1 and S2.

### Immunocytochemistry

Cells were fixed with 4% paraformaldehyde (PFA) in PBS for 45 min at 4°C and permeabilized with 0.5% Triton X-100 in PBS for 30 min at room temperature. Samples were treated with the blocking solution A (0.1%BSA, 0.2% TritonX-100, and 0.05% Tween20 in PBS), for 2 h at room temperature and subsequently treated with a blocking solution A with 1%BSA overnight at 4°C. Consequently, they were incubated with an antibody against SM22α (1:200, ab14106, Abcam, Cambridge, UK) in the blocking solution A overnight at 4°C. After washed several times with solution A, cells were incubated overnight at 4°C and for 2 h at room temperature in solution A containing Goat anti-Mouse IgG (H+L) Cross-Adsorbed Secondary Antibody, Alexa Fluor 555 (1:200, A-21422, Invitrogen). F-actin was stained with Alexa Fluor 488-conjugated phalloidin (1:200, A12379, Invitrogen) by incubating cells overnight at 4°C. Nuclei were stained with Hoechst 33342 (1:1000, H3570, Invitrogen) by incubating cells overnight at 4°C. The samples were stored at 4°C until observation with a microscope.

### Microscopy

To visualize cell morphology under a confocal microscope, HUVECs were labeled with 4 µM CellTracker^TM^ Violet BMQC (Thermo Fisher Scientific) or CellTracker^TM^ Green CMFDA (Thermo Fisher Scientific) in EGM-2 in a 60-mm dish before the fabrication of the semi-3D or 3D coculture model. Microvessels were stained with 2 mg/mL Fluorescein Ulex Europaeus Agglutinin 1 (UEA1, Vector Laboratories, Burlingame, CA, USA) in EGM-2 daily until microscope observation. For time-lapse observation, bright-field and fluorescent images were captured using a fluorescent microscope (Axio observer Z1) equipped with a 20×-observation lens. Z-stack images were taken using CLSM equipped with a 20×-objective lens at 30-min or 1-h intervals. Red, green, and blue fluorescence were detected using laser wavelengths of 555, 488, and 405 nm, respectively. The acquired Z-stack images were exported from the IMARIS software (version 9.0.0, BitPlane, Zurich, Switzerland).

Bright-field and fluorescent images were captured using a fluorescent microscope (Axio observer Z1) equipped with a 20×-observation lens. Z-stack images were taken using CLSM equipped with a 40× water-immersion detection objective lens. Red fluorescence, green fluorescence, and Hoechst 33342 were detected using laser wavelengths of 555, 488, and 405 nm, respectively. The images were processed using a median filter (3×3×3), and the maximum intensity projection (MIP) images were obtained with the ZEN 2 blue edition software (Carl Zeiss). For the observation of the semi-3D coculture model, samples were placed upside and down, and the glass-bottom reservoir filled with PBS.

### Image analyses

#### Quantification of morphological changes in the microvessel layer in 3D microvessel-on-a-chip

The Z1 images of the tumor–microvessel models at day 5 were used. The edge of the microvessel layer nearby tumor organoids, 500 µm from the nearest point from organoids, were defined as the region of interest (ROI). *Invasion on microvessel* was calculated by dividing the total ROIs hijacked by tumor organoids in the NI models by total ROIs of all samples observed for each culture condition. *Microvessel layer disruption* was calculated by dividing the number of the total disrupted ROIs in the DI models by total ROIs observed of all samples for each culture condition.

#### Quantification of invadopodium formation

The Z-stack images of tumor organoids in the DI models at day 3 were used. 3D images were reconstructed from Z-stack images and invadopodia were manually measured using the IMARIS software (version 9.0.0). Organoids with at least 1 invadopodium (length >20 µm) were defined as invadopodium-positive organoids, and the invadopodium-positive ratio was calculated by dividing the total number of invadopodium-positive organoids by all counted organoids of all samples for each cell type. The Z1 images of cancer organoids in the semi-3D models at day 1, 3, and 6 were used for the temporal change in cancer invadopodium formation. The number of invadopodia was manually counted, and the number of invadopodia per 1 organoid was calculated by dividing the total number of invadopodia by the number of organoids in each sample at the same day for each culture condition.

### Statistical analysis

The data were analyzed using two-tailed unpaired Student’s *t*-tests, Chi-square, Fisher’s exact tests, one-way ANOVA with Bonferroni’s multiple comparisons test or two-way ANOVA with Šídák’s multiple comparisons test using Graphpad Prism10.0.2 (GraphPad Software, Boston, MA, USA) Values are presented as mean ± standard deviation (s.d.). Differences between means were considered statistically significant at *p* < 0.05. All data were reproduced with at least two independent experiments.

## Acknowledgments

We thank Dr. Hitoshi Niwa (Kumamoto University, Japan) for providing the pPB-CAG-IP PiggyBac transposon expression vector, Dr. Kazuo Shinya (AIST, Japan) for the technical advice on the size-controlled organoid formation, Dr. Mikako Shirouzu (RIKEN, Japan) and Dr. Takehisa Matsumoto (RIKEN) for the preparation of Fc-chimeric proteins, Joris Pauty (The University of Tokyo, Japan) for the design of the 3D culture platform, and Dr. Katarzyna A. Podyma-Inoue (Tokyo Medical and Dental University) for the technical advice on the Fc-chimeric protein experiment.

## Funding

AMED P-CREATE JP21cm0106272 (Y.T.M.)

AMED P-CREATE JP22ama221205 (T.W.) the Extramural Collaborative Research Grant of Cancer Research Institute, Kanazawa University (H.O., Y.T.M.)

Grant-in-Aid for Scientific Research (B) JP20H03851 (T.W.)

JSPS KAKENHI Grant-in-Aid for JSPS Fellows JP23KJ0490 (Y.I.) the Sasakawa Scientific Research Grant from The Japan Science Society (Y.I.)

WINGS-QSTEP (Y.I.)

## Author contributions

Conceptualization: M.O. and Y.T.M.

Design of the experiments: Y.I., H.O., J.S., M.O., and Y.T.M.

Writing the paper: Y.I., J.S., H.O., M.O., and Y.T.M.

Fabrication of the device and characterization of samples: Y.I.

Performing experiments using the device (3D or semi-3D model): Y.I.

Performing the organoid transplantation experiments: S.Y.K. and H.O.

Performing the qPCR experiments: Y.I. and J.S.

Primer design: H.S.

Design of inhibition experiments: K.T. and T.W.

Supervision: Y.T.M. and M.O.

Discussion: Y.I., J.S., H.O., S.Y.K, K.T., H.S., T.W., M.O., and Y.T.M.

## Data and materials availability

All data are available in the main text or the supplementary materials.

## Supplementary Materials

### Summary

Table S1. List of primer pairs for RT-qPCR.

Table S2. List of primer pairs for the quantification of RNA copies.

Movie S1 (.mp4 format). Time-lapse imaging of AKTP near-interaction model for the detection of vessel co-option and the release of CTC clusters from AKTP organoids.

Movie S2 (.mp4 format). Time-lapse imaging of AP near-interaction model.

Movie S3 (.mp4 format). Time-lapse imaging of AKTP distant-interaction model for the detection of AKTP invasion and partial endothelial cell migration.

Movie S4 (.mp4 format). Time-lapse imaging of AP distant-interaction model.

