## Supplementary file for "Tumor-microvessel on-a-chip reveals sequential intravasation cascade of cancer cell clusters"

**This PDF file includes:**

- Table S1. List of primer pairs for RT-qPCR
- Table S2. List of primer pairs for the quantification of RNA copies
- Legends for movies S1 to S4

**Other Supplementary Materials for this manuscript include the following:**

- Movie S1 (.mp4 format). Time-lapse imaging of AKTP near-interaction model for the detection of vessel co-option and the release of CTC clusters from AKTP organoids.
- Movie S2 (.mp4 format). Time-lapse imaging of AP near-interaction model.
- Movie S3 (.mp4 format). Time-lapse imaging of AKTP distant-interaction model for the detection of AKTP invasion and partial endothelial cell migration.
- Movie S4 (.mp4 format). Time-lapse imaging of AP distant-interaction model.

**Supplementary tables:**

**Table S1. List of primer pairs for RT-qPCR**

| Target | Primer pairs for qPCR |
|---|---|
| **mouse *Tgfb1*-f** | ATTCAGCGCTCACTGCTCTTGT |
| **mouse *Tgfb1*-r** | TTCCAACCCAGGTCCTTCCTAA |
| **mouse *Tgfb2*-f** | ATAATTGCTGCCTTCGCCCT |
| **mouse *Tgfb2*-r** | CCCCAGCACAGAAGTTAGCATT |
| **mouse *Inhba*-f** | GGGGAGAACGGGTATGTGGA |
| **mouse *Inhba* -r** | CCTGACTCGGCAAAGGTGAT |
| **human *TGFB1*-f** | GGAAATTGAGGGCTTTCGCC |
| **human *TGFB1*-r** | CCGGTAGTGAACCCGTTGAT |
| **human *TGFB2*-f** | GTTCGATTTGACGTCTCAGCAAT |
| **human *TGFB2*-r** | CAATCCGTTGTTCAGGCACTCT |
| **human *INHBA*-f** | CCTCCCAAAGGATGTACCCAA |
| **human *INHBA*-r** | CTCTATCTCCACATACCCGTTCT |
| **human *TAGLN*-f** | TCAAGCAGATGGAGCAGGTG |
| **human *TAGLN*-r** | GCTGCCATGTCTTTGCCTTC |
| **human *ACTA2*-f** | CAAAGCCGGCCTTACAGAG |
| **human *ACTA2*-r** | AGCCCAGCCAAGCACTG |
| **human *FN1*-f** | AAACCAATTCTTGGAGCAGG |
| **human *FN1* -r** | CCATAAAGGGCAACCAAGAG |
| **human *KDR*-f** | CAGAATCCCTGCGAAGTACCTT |
| **human *KDR* -r** | GTCAGTACATGCCCCGCTTTAA |
| **human *TEK*-f** | GGTGGAAAAGCCCTTCAACA |
| **human *TEK*-r** | CATCCCCAAAGTAAGGCTCAG |
| **human *PPIA*-f** | TGGTTCCCAAGTTTTTCATCTGC |
| **human *PPIA*-r** | CCATGGCCTCCACAATATTCA |
| **mouse *Ppia*-f** | GAGCTGTTTGCAGACAAAGTTC |
| **mouse *Ppia*-r** | CCCTGGCACATGAATCCTGG |

**Table S2. List of primer pairs for the quantification of RNA copies**

| Target | Primer pairs |
|---|---|
| **mouse *Inhba*-f** | CCCTAGTGTTAAAGTGGCTC |
| **mouse *Inhba* -r** | AAAGTGTCAATGAAGCTTACAAG |
| **human *INHBA*-f** | AGGTGGGTGTGGTGAGAAAA |
| **human *INHBA* -r** | CACACTGTTTCTGCAGGTTCC |
| **mouse *Tgfb1*-f** | CTTGCAGAGATTAAAATCAAGTGTG |
| **mouse *Tgfb1*-r** | TCAGGCGTATCAGTGGGG |
| **human *TGFB1*-f** | GAGGACTGCGGATCTCTGTG |
| **human *TGFB1*-r** | GCACTTCAACAGTGCCCAAG |

### Supplementary movies:

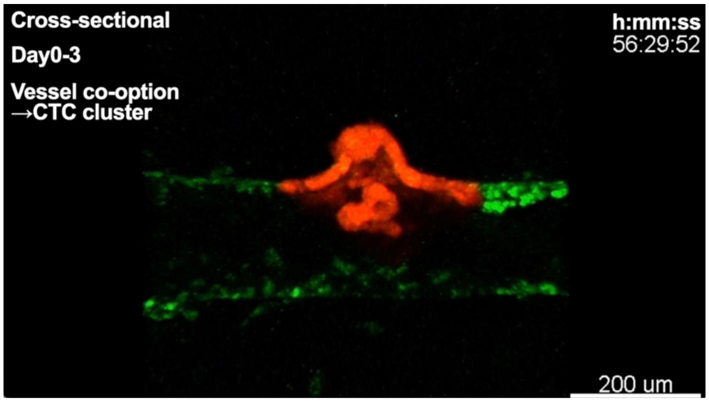


**Movie S1. Time-lapse imaging of AKTP near-interaction model for the detection of vessel co-option and the release of CTC clusters from AKTP organoids**. AKTP organoids were labeled with tdTomato (red), and the microvessels are stained with UEA-1 (green).

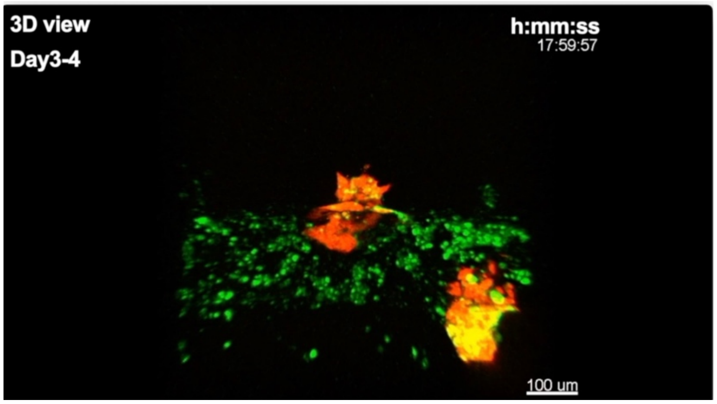


**Movie S2. Time-lapse imaging of AP near-interaction model.** AP organoids were labeled with tdTomato (red), and the microvessels are stained with UEA-1 (green).

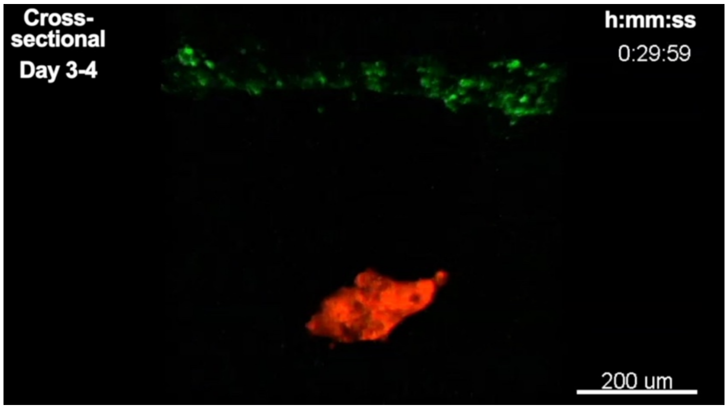


**Movie S3. Time-lapse imaging of AKTP distant-interaction model for the detection of AKTP invasion and partial endothelial cell migration.** AKTP organoids were labeled with tdTomato (red), and the microvessels are stained with UEA-1 (green).

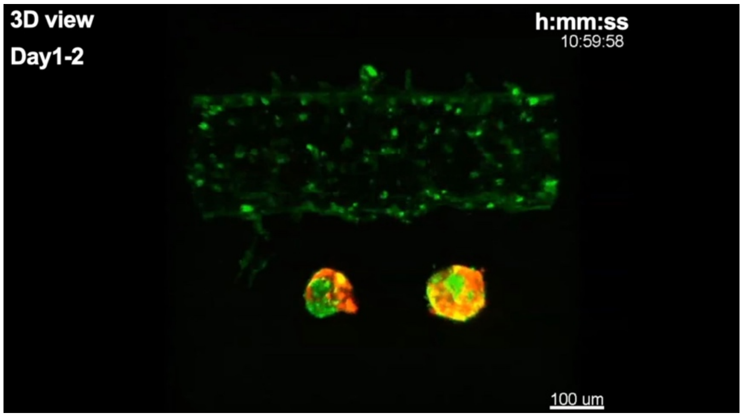


**Movie S4. Time-lapse imaging of AP distant-interaction model.** AP organoids were labeled with tdTomato (red), and the microvessels are stained with UEA-1 (green).
